# Better expression profile of CD8 and CD4 gene in uterus of pregnant ewe in comparison to non-pregnant - a novel report

**DOI:** 10.1101/2020.10.07.329730

**Authors:** Md Mofijul Islam, Aruna Pal, Partha Das, Samiddha Banerjee

## Abstract

CD8 and CD4 T cells play a central role in the immune response to viruses and intracellular pathogens as well as functions for the maintenance of both the mother and fetus. The present study was conducted to explore the differential gene expression profile for CD8 and CD4 present in the uterus with reference to the gravid and non-gravid Garole sheep and confirmation through immuno histochemical studies. Better CD8 and CD4 gene expression was observed in the mid uterus of pregnant ewes compared to that of non-pregnant ewes, where CD8 expression was better to that of CD4. Gene expression profiling of CD8 and CD4 are reported here for the first time in sheep. CD8 and CD4 expression may be regarded as the useful factor for maintenance of pregnancy. The current observations demonstrate that during pregnancy in ewe the immune system may respond to changes in the maternal environment to maintain the size and function of the CD8 and CD4 T-cell compartment. CD8 and CD4 expression may be employed as marker for pregnancy detection in sheep, which remains always a challenge for sheep breeders.

## Introduction

Abortion or preterm pregnancy is one of the major problem in both human and livestock sector. Lymphocytes play key roles in the defense of the body. T-lymphocytes regulate acquired immunity and are responsible for cell mediated immune response. CD8+ T-cell is a T-lymphocyte also known as killer cell or cytotoxic T cell and they act as class-I MHC receptor molecules. It is very important for immune defense against intracellular pathogens, including viruses and bacteria. CD8+ T cells serve as important mediator for fetal tolerance and antiviral immunity. Cytotoxic T cells act through T-cell receptors (TCRs) recognizing a specific antigen. Antigens within the cell are attached to class I MHC molecules and in the surface of the cell, they are recognized by the T cell and destroyed.^**1**^

CD8 T cells play a central role in the immune response to viruses and intracellular pathogens, so the maintenance of both the number and function of these cells are critical to protect both the mother and fetus. The proportion of maternal CD8 T cells in the spleen and the uterine draining lymph nodes are gradually increased at mid-gestation, which correlates with enhanced CD8 T-cell proliferation and an increased relative expression of both pro-survival and pro-apoptotic molecules.^**2**^

CD4 and CD8 T cells are surface molecules play a role in T cell recognition^**3**^ and they are activated by binding to the MHC-II and MHC-I ligands respectively on an antigen presenting cell (APC)^**4**^. CD4 cells are also known as T helper cells because they produce various cytokines required for the activation of leukocytes and send signals to other types of immune cells, including CD8 killer cells, which then destroy the infectious particle. CD4 T cells secrete IFN-γ that maximizing bactericidal activity of phagocytes such as macrophages and digest intracellular bacteria^**5**^ and protozoa^**6**^. Sometime these cells suppress immune responses. If CD4 cells become depleted, for example in untreated HIV infection, the body is left vulnerable to a wide range of infections^**7, 8**^. Understanding exactly how T helper cells respond to immune challenges is currently of major interest in immunology, because such knowledge may be very useful in the treatment of disease and in increasing the effectiveness of vaccination.

Garole sheep (*Ovis aries*) are distributed in the Sundarban region of West Bengal in India. It is known for its prolificacy and adaptation to the saline marshy land. It is believed that these sheep contributed to the prolificacy gene (Boorola fecundity) in Merino sheep of Australia. Flocks are stationary and average flock size ranges from three to five. Twin and triplet births are common. Average adult weights for male and female Garole was observed to be 10.3 ± 0.31 kg 9.72± 0.28 kg, respectively^**9**^. They are reared for mutton production ^**10**^. Since Garole sheep possess the unique characteristic for reproduction, Garole sheep was choosen as model organism for the current study. Earlier we had studied the female reproduction of Ghungroo pig using mitogenetics approach^**11**^.

Successful pregnancy requires the maternal immune system to tolerate the semi-allogenic fetus. A failure in immune tolerance may result in abnormal pregnancies, such as recurrent spontaneous abortion and it happens due to several etiological factors. Out of this, immune deficiency causes several infections in the female reproductive system which results in misconception or abortion. The most notable infectious diseases of the reproductive tract of sheep are those that infect the placenta and cause abortion. These infections outreach when the natural immune barrier broken down^**12**^.

Considering the above facts, the current study was conducted to explore the differential mRNA expression profile and immunohistochemical studies for the identification of immune cells (Lymphocytes) as CD8 and CD4 expression in both the gravid and non-gravid Garole sheep.

## Results

### Gene expression profiling with real time PCR

We analysed gene expression of CD8 and CD4 mRNA, normalized for 18S rRNA expression and was quantified by using SYBR green PCR system from samples (Tip and middle horn of the uterus) of gravid and non-gravid ewes.

Out of the two different parts of the uterus, the middle horn of the uterus revealed better expression of both CD8 (Fig 1A) and CD4 (Fig 1B) in comparison to the tip of Horn of the uterus.

**Fig 1.**
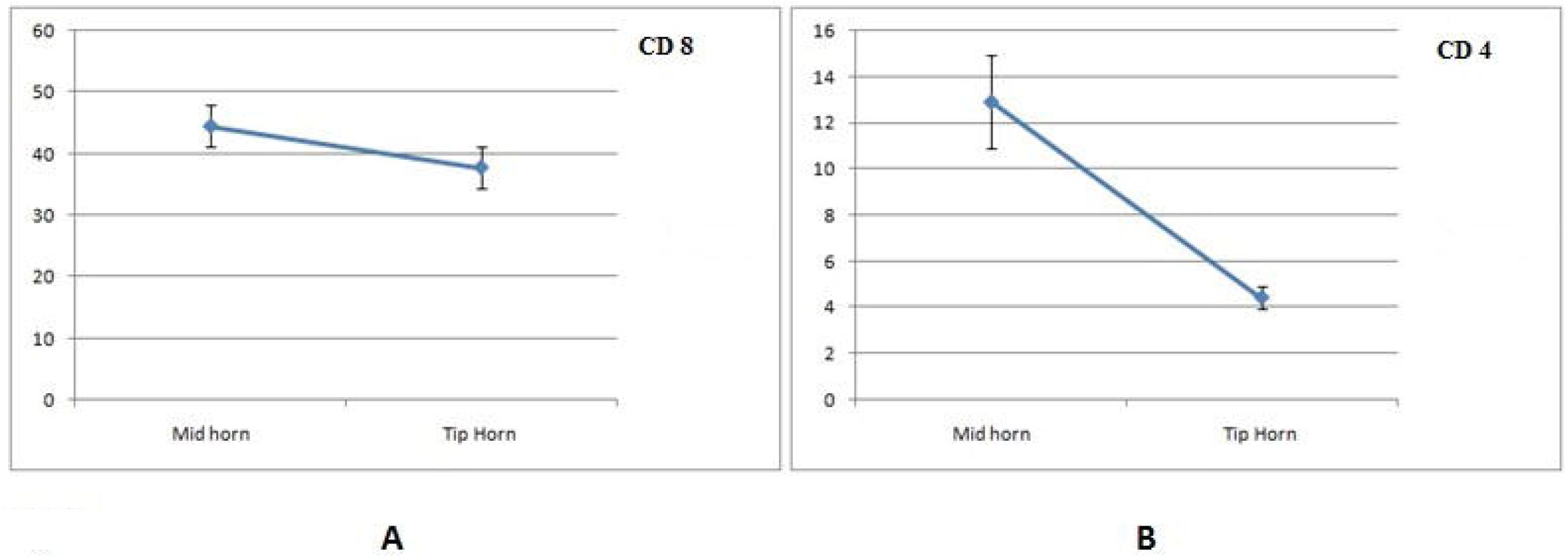
A. Differential mRNA expression profile of CD8 in middle and tip of the horn of uterus of ewe B. Differential mRNA expression profile of CD4 in middle and tip of the horn of uterus of ewe (Middle of the horn had maximum expression compared with tip of the horn)

An increased expression of both CD8 (Fig 2A) and CD4 (Fig 2B) mRNA in gravid uterus (pregnant) as compared to the non-gravid (non-pregnant) uterus was reported. The current observations demonstrate that during pregnancy in ewe the immune system responds to changes in the maternal environment to maintain the size and function of the CD8 T-cell compartment.

**Fig 2.**
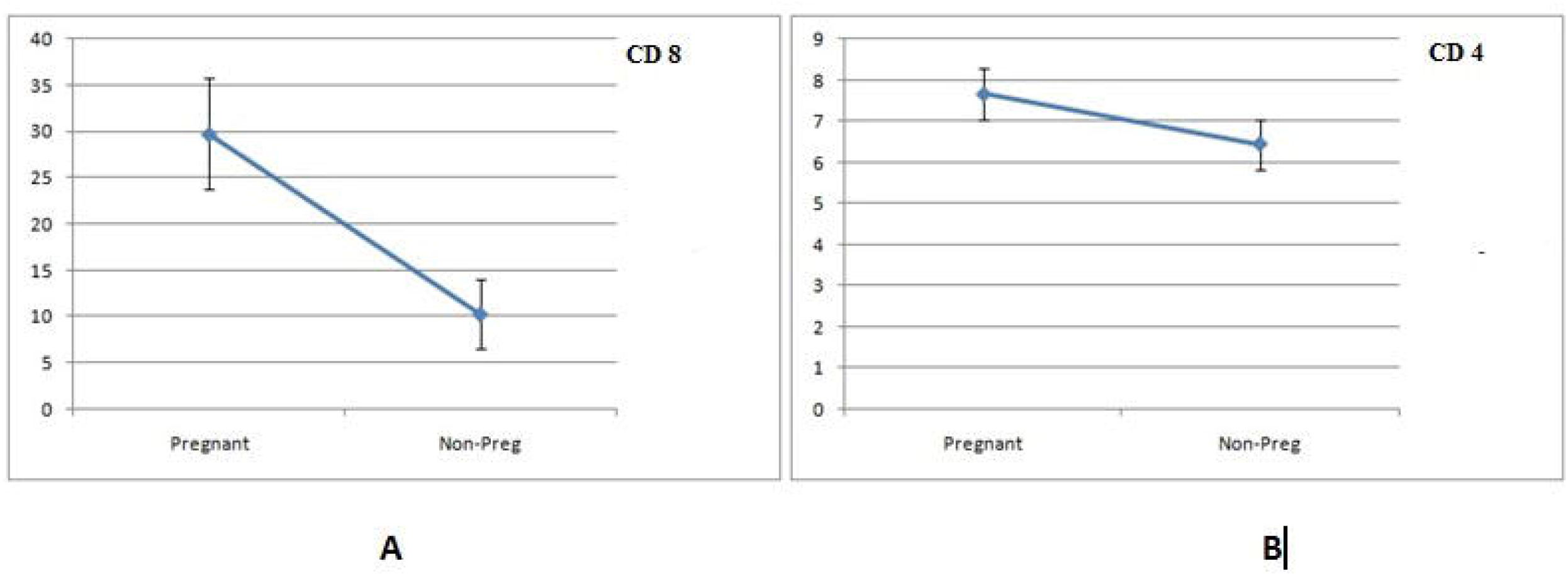
A. Differential mRNA expression profile of CD8 with respect to Pregnant and Non-pregnant uterus of ewe. B. Differential mRNA expression profile of CD4 with respect to Pregnant and Non-pregnant uterus of ewe. (Gravid or pregnant uterus had increased expression than non-gravid uterus)

It has also been observed that CD8 expression was higher in comparison to CD4 expression in both the cases of middle and tip of the horn of the uterus (Fig 3A). The similar phenomenon of better expression profile of CD8 compared to that of CD4 was also observed in case of both pregnant and non-pregnant uterus of ewe (Fig 3B)

**Fig 3.**
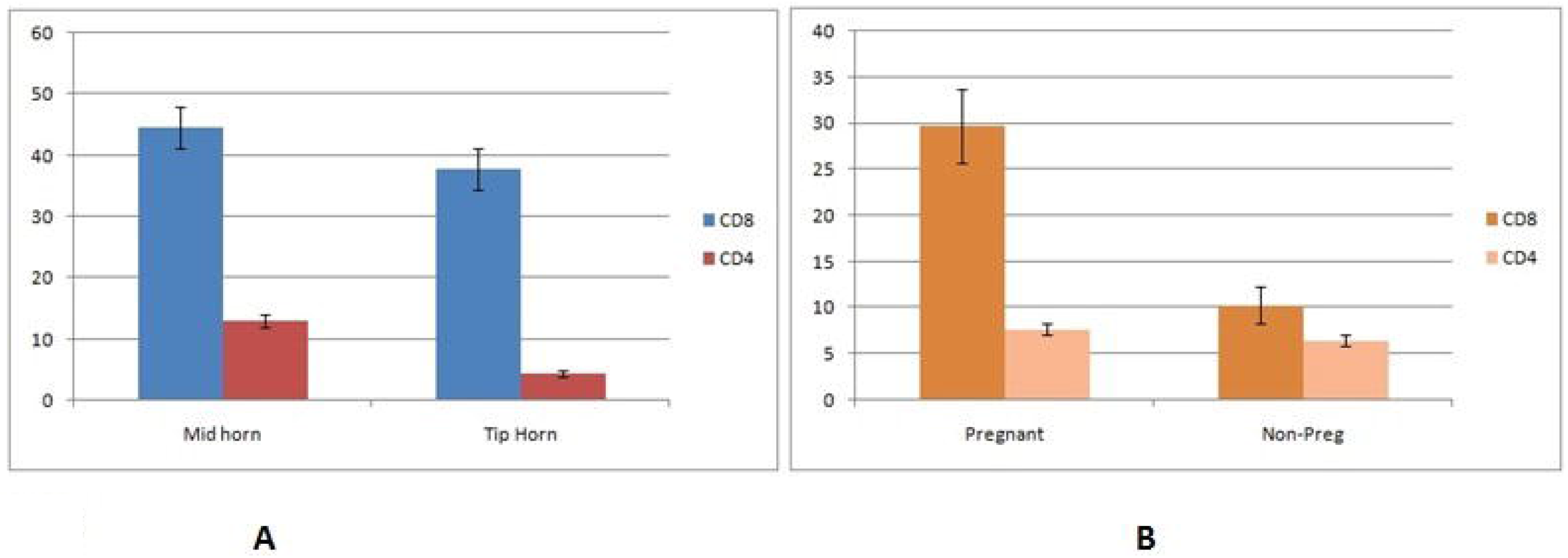
A. Differential mRNA expression level of CD8 and CD4 in different parts of uterus of ewe-in Middle and Tip of the horn. B. Differential mRNA expression level of CD8 and CD4 in uterus of pregnant and non-pregnant ewes.

### Immunohistochemistry

The distribution of lymphocytes were identified by routine H & E staining of the tissues with a characteristics of a dark staining round nucleus that occupied virtually the entire cells. Lymphocytes were found in all the tissue samples under investigation. Lymphocytes were not equally distributed in the different parts of uterus and they were especially abundant in the basal zone of endometrium. Irrespective of the stage of the physiological status of the uterus, lymphocytes were found to infiltrate the caruncular and inter caruncular epithelium Lymphocyte population was also identified in the sub epithelial area, the stroma. Lymphocytic aggregation was however found in none of the tissue irrespective of physiological condition. The epithelium of the endometrial glands and their ducts were frequently infiltrated by lymphocytes. However, the lymphocyte population was comparatively more at the middle horn of the uterus as compared to the tip and body of the uterus in both gravid and non-gravid ewes.

Immunohistochemical study evidenced the presence of cells displaying immunoreactivity for CD8+ antibody, in the luminal, glandular epithelial layers and stroma of the uterus of both gravid and non-gravid ewes.

In gravid uterus of ewe, the CD8+ lymphocyte sub population were more at the middle of the horn (Fig 4A) compared to the tip of the horn (Fig 4B). The occurrence of CD8+ cells were found within the luminal and glandular epithelium of the caruncular and inter caruncular areas. The CD8 population was observed to be better in the stroma compared to that of the epithelium. In some areas within the caruncular stroma, positive cells appeared in the form of clusters. Similarly, CD4 cells were observed to be better expressed in middle of the horn (Fig 5A) in comparison to that of tip of the horn (Fig 5B).

**Fig 4.**
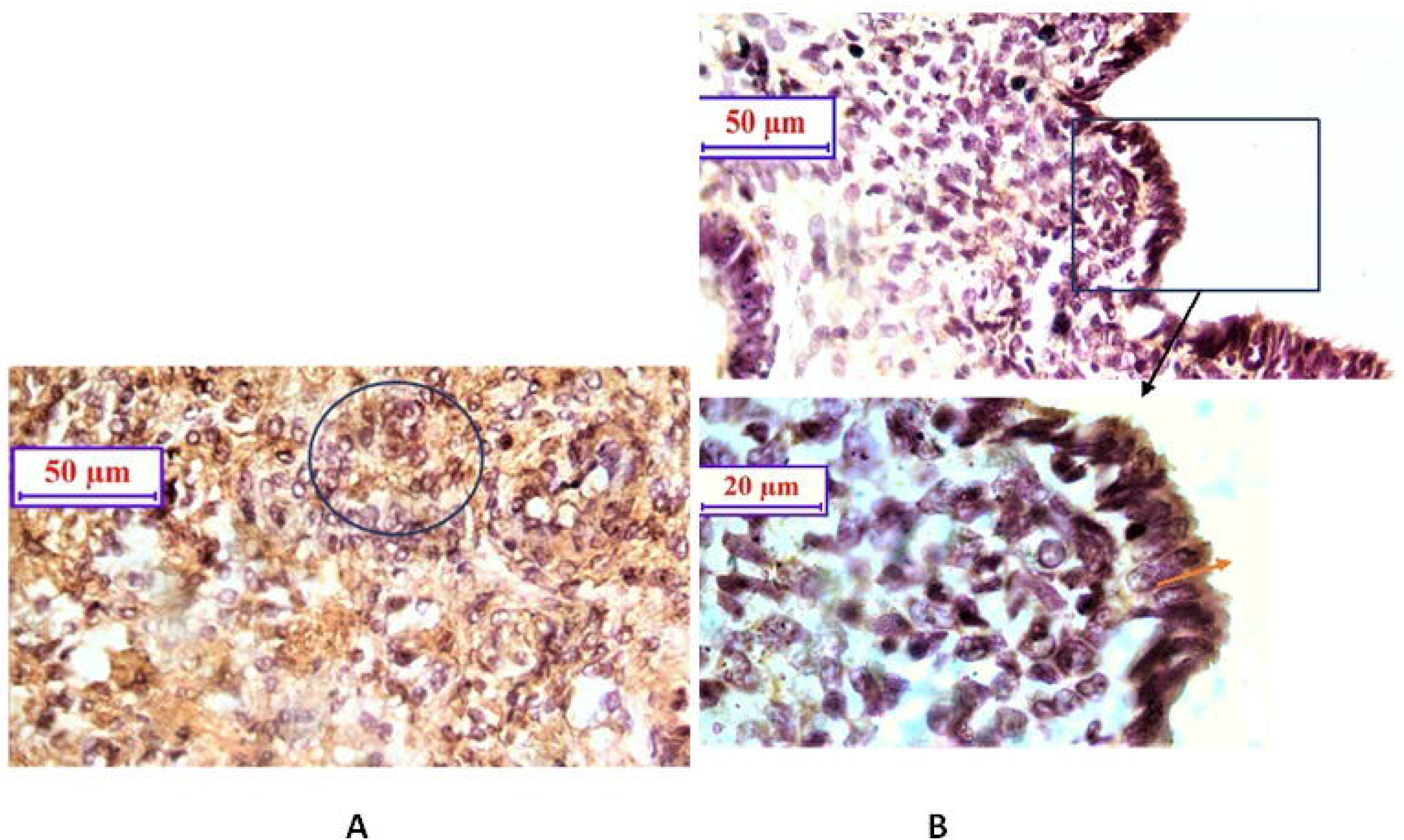
A. Photomicrograph of Immunohistochemistry showing CD4 lymphocytes distributed in stromal area of ***middle horn*** of gravid ewe in a cluster form (circle area). IHC 40X. B. Photomicrograph of Immunohistochemistry showing CD8 lymphocytes localisation at the stroma of inter caruncular area in ***tip of the horn*** of gravid ewe. CD8 positive lymphocytes (arrows). IHC 40X (above) & 100X (below).

**Fig 5.**
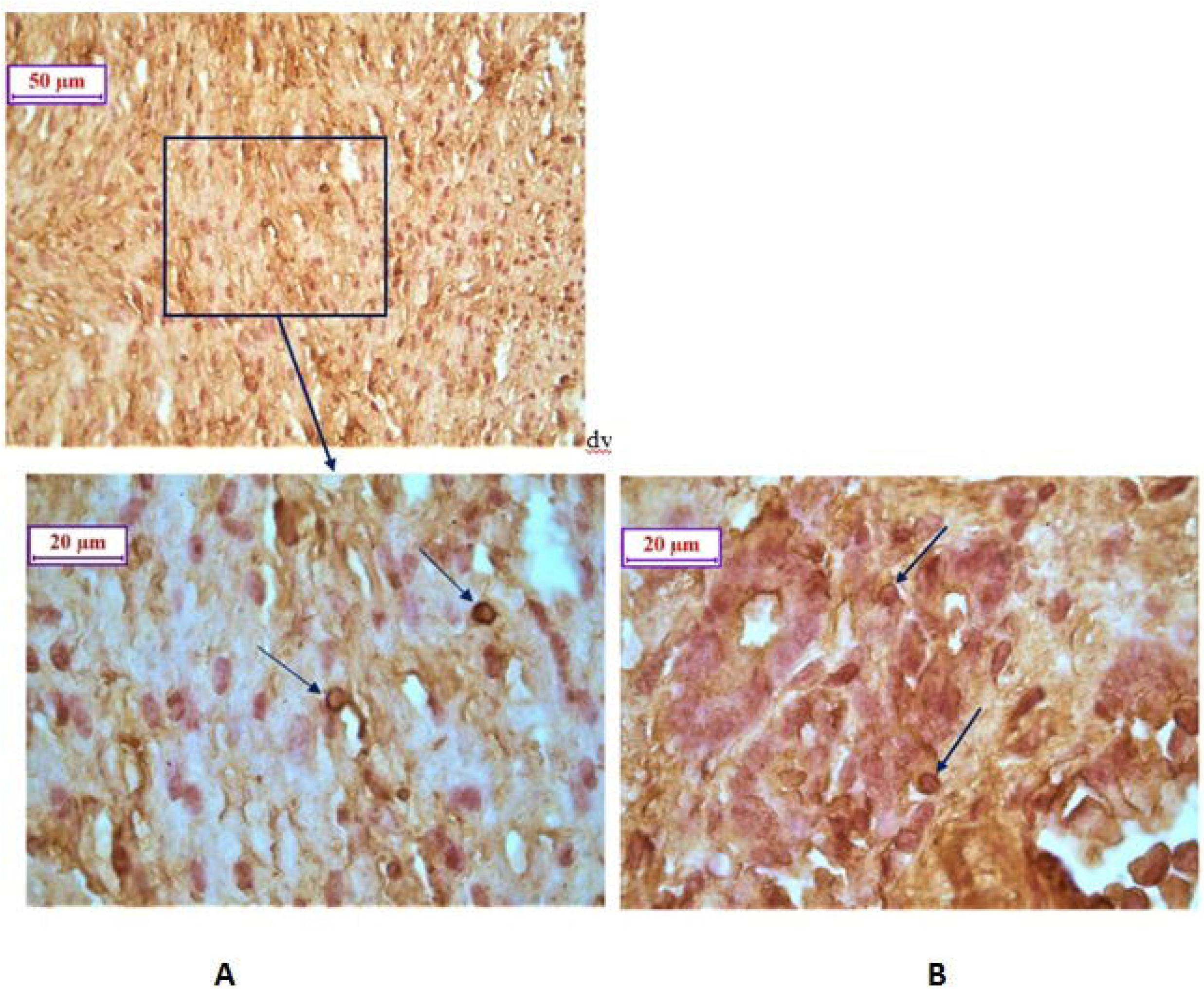
A. Photomicrograph of Immunohistochemistry showing CD4 positive cells (arrows) interspersed at the stromal region of caruncle of ***middle horn*** of uterus of gravid ewe. T. Serosa (S), T. Muscularis (M) and Caruncles (C). IHC 40X & 100X. B. Photomicrograph of Immunohistochemistry showing CD4 positive cells (arrows) distributed at the stromal region of caruncle of ***tip of the horn*** of uterus of gravid ewe. IHC 100X

In gravid uterus (Fig 6A) the population of CD8+ cell was observed to be better in comparison to that of non-gravid uterus (Fig 6B). Pregnancy was associated with an increase in the thickening of the epithelial lining of the endometrial glands and lumen but the distribution of lymphocyte was unaffected by pregnancy status. However the number of CD8+ cells was more in stromal region compared to luminal epithelium. CD4+ cells were also observed to be distributed mostly in the stroma and sub-epithelial region of both the caruncular and inter caruncular area of uterus like CD8 cells. CD4 expression was also observed to be better in the uterus of pregnant ewe(Fig 7A) compared to that of non-pregnant ewe (Fig 7B).

**Fig 6.**
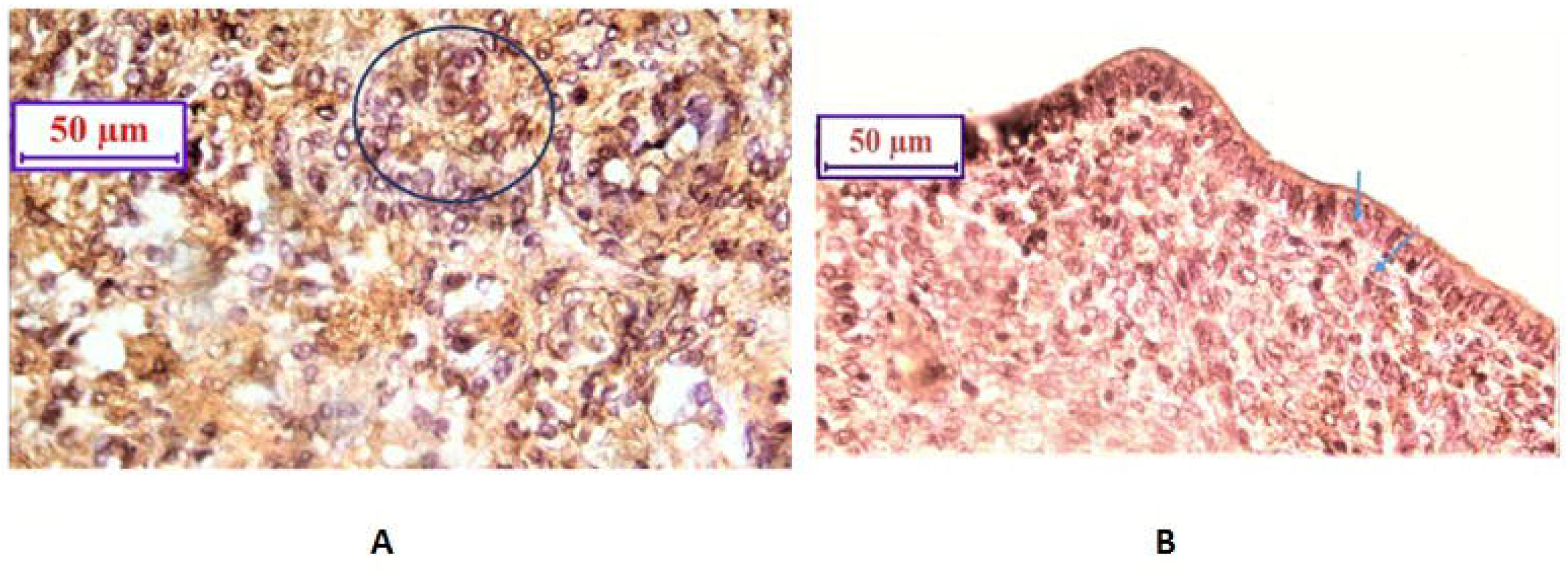
A. Photomicrograph of Immunohistochemistry showing CD8 lymphocytes distributed in stromal area of caruncular region of the middle horn of ***gravid*** ewe in a cluster form (circle area). IHC 40X. B. Photomicrograph of Immunohistochemistry showing CD8 lymphocytes, interspersed at the stromal region of caruncular area of middle horn of uterus of non-gravid ewe. CD8 positive lymphocytes (arrows) at sub epithelial region. IHC 40X.

**Fig 7.**
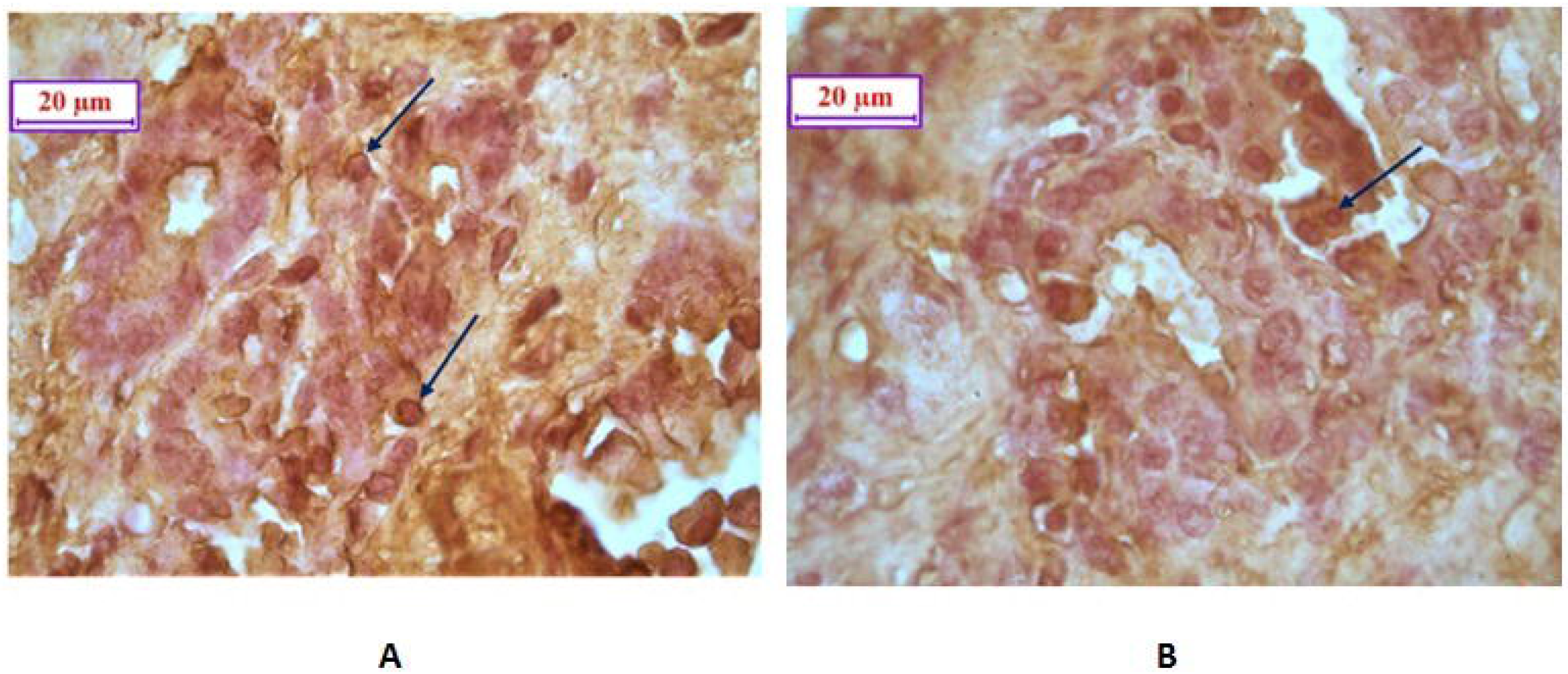
A. Photomicrograph of Immunohistochemistry showing CD4 positive cells (arrows) distributed at the stromal region of caruncle of tip of the horn of uterus of gravid ewe. IHC 100X. B. Photomicrograph of Immunohistochemistry showing CD4 positive cells (arrow) interspersed at the stromal region of caruncle of tip of the horn of uterus of non-gravid ewe.IHC 100X.

Comparison of CD8 and CD4 has revealed some interesting findings. In gravid uterus, the expression of CD8 was observed to be higher compared to that of CD4 gene expression.

## Discussion

**CD8** (cluster of differentiation 8) is a transmembrane glycoprotein that serves as a co-receptor for the T cell receptor (TCR). **CD8** binds to a specific class I MHC protein (major histocompatibility complex) molecules. Endometrial mucosa contains a set of immune cells, which apart from providing host-pathogen immunity, have immense role in implantation and maintenance of pregnancy. The mucosal surface of the uterus (endometrium) contains both innate and adaptive immune cells. The implanted foetus has different genetic makeup from its mother and simultaneously differ in immunological makeup to that of the mother. Endometrial mucosal cells of the mother tolerate the semi allogenic foetus without loss of host immunity^**13**^. The peculiarity of endometrial CD8-T cells has memory CD8-T cells. Memory CD8 T –cells were less in the patients with recurrent miscarriage^14,15^.

Abortion or tissue rejection is one of the major problems in vogue. Tissue rejection is generally mediated by T lymphocytes. In the current study, it was observed that CD8 was genetically expressed more in pregnant uterus compared to non-pregnant. It was also observed that CD8 was highly expressed in middle of the horn compared to that of tip of the horn which clearly indicated that CD8 expression is essential for maintenance of pregnancy. Similar results were also observed by other researchers. The greatest number of T-lymphocytes in the sheep endometrium was observed in epithelial layer of the glandular and luminal epithelium or in stroma immediately adjacent to the epithelium^**16, 17**^. The near absence of stromal T-lymphocyte may indicate that blood was not a major source of endometrial T-lymphocytes. In fact many endometrial inter epithelial lymphocytes developed extra thymically rather than being derived from a circulating pool of lymphocytes^**18**^.

In sheep, cytotoxic (CD8+) T cells were reported to be located primarily adjacent to the epithelial layer of the uterus ^**19, 20**^ and similar observation in sow were reported^**21**^. The presence of cells displaying immunoreactivity for both CD 4+ and CD 8+ antibodies in the epithelial layer and stroma of the cervix and uterine horn of female goats were reported^**22**^. However, only an occasional CD8+ cell was observed to be present in the caruncular and intercaruncular stroma and the loose connective tissue in the vicinity of the endometrial glands in sheep^**17**^. It was identified that in most sections, CD4+ and CD8+ cells were rarely observed or absent^**16**^, although for a few ewes positively staining cells were readily apparent. This was not in accordance with our present findings.

In non-gravid uterus the CD8+ lymphocyte sub population was found to be more at the middle of the horn. However, the CD8+ cells increased markedly during the mid-to-late luteal and follicular phases, respectively, of the cycle^**20**^.

In gravid uterus the population of CD8+ cell was found to be comparatively greater. There was a tendency for higher number of CD8+ cells in luminal epithelium in pregnant ewes^**23**^ and there was no significant effect of uterine type on number of CD8+ cells in either luminal or glandular epithelium. The role of immune competent cells was documented^**24**^ who observed that acute inflammation in the pig endometrium in response to fertile mating which included marked changes in the tissue and immune cell components of the endometrium. There appeared to be suppression and activation of various immune cell components in the uteri of pregnant pigs. This phenomenon was presumably in response to foetal or trophoblast antigens, suggesting that the local immune system was involved in the uterine response to pregnancy. The population was very less at the palcentomal region of the endometrium and also in the inter placentomal endometrium^**24,25**^.

Leucocytes were a significant population in the human endometrium (EM), accounting for 10–20% of all endometrial cells. Of the leucocyte population, ≈50% were T cells and at least two-thirds of these were CD8^+^ T cells. These CD8^+^ cells were found in three distinct locations within the EM: as scattered stromal cells; as intra epithelial lymphocytes; and in lymphoid aggregates (LA). LA develop in the stratum basalis of the human EM during the proliferative phase of the menstrual cycle, but were absent in the EM of post menopausal women, suggesting that their development was hormonally influenced^**26**^.

In sheep very few lymphocytes carrying classical B and T cell markers (CD5, surface immunoglobulin) were detected in the uterine epithelial cell suspensions and all intra epithelial lymphocytes (IEL) expressed the CD8 surface marker although with varying intensities^**21**^, which was in support of the present findings. It was revealed that MHC-I and - II^+^ cells were expressed by individual cells in organ layers, while CD8+ T cells and CD68+ macrophages were dominant in epithelium and muscle layer of bovine^**27**^.

Natural killer (NK) cells were observed to be essential for establishment of human and rodent pregnancies. The function of these and other cytotoxic T cells (CTL) was controlled by stimulatory and inhibitory signalling^**28**^. An increased proportion of NKp46+ and CD8+ cells were observed in the uterus of pregnant heifers, which supports our observation.

During human pregnancy, the tolerance of the semi-allogeneic fetus presents a significant challenge to the maternal immune system. In a woman, who had previous pregnancy, T cells were detected with specificity for foetal epitopes. Although it was previously thought that for maintaining maternal tolerance of the foetus, such foetal specific cell were generally deleted during pregnancy. Fetal-specific CD8+ T lymphocytes were observed in half of all pregnancies and expression increased from the first trimester upto the postnatal period. Fetal-specific cells lead to an effector memory phenotype and were broadly functional. These retain their ability for proliferation, secrete IFN-γ, and lyse target cells and recognition of naturally processed peptide on male fetal cells. Hence, fetal-specific adaptive cellular immune response is a normal consequence of human pregnancy, fetal-specific T cells are not deleted during human pregnancy^**29**^.

Treg (the naturally arising regulatory cells) were found to be essential for surface expression of CD4 and CD25 (IR2Rα chain)^**30**^. In mouse model, it has been studied that fetal/ placental antigen presentation and recognition of fetal antigen are mediated by CD4^+^ T cells through indirect recognition. In human pregnancy, there is evidence of HLA-C mismatch leading to an increase of CD4^+^CD25^dim^ activated T cells and the presence of functional CD4^+^CD25^bright^ regulatory T cells.^**31**^ In human pregnancy also, similar results were obtained i.e. indirect recognition of HLAC allele.

The mammalian uterus is unique in that it possesses the ability to mount an immune response against pathogenic organisms while not usually responding to ‘foreign’ antigens on sperm and fetal allografts^**32**^. Mucosal sites inhabited by commensal flora which are responsible to generate protective immune responses in the lower female reproductive tract, whereas the upper reproductive tract is normally sterile^**33**^.

The uterus contains populations of T cells sufficient to induce graft rejection^**34,35**^. Proper adjustments in uterine lymphocyte function to inhibit anti-fetal responses are therefore critical for successful pregnancy and disruption of this regulation can lead to pregnancy loss^**36**^.

Studies were conducted at genomic level through differential gene expression study of CD8 and CD4 gene. It was confirmed that both CD8 and CD4 gene expression was more in pregnant compared to that of non-pregnant uterus. Gene expression was also observed to be better in middle of the horn compared to that of tip of the horn for both genes. Since in sheep, implantation occurs in the horn, it is evident to have better CD8 and CD4gene expression. This is the first report to study the expression level of CD8 in sheep, hence comparison was not possible. The study was further confirmed with immune histochemical studies which showed the reactivity of CD8 cells in gravid and non gravid ewes. This is the first report of CD4 gene expression in sheep with respect to pregnancy. However some preliminary studies revealed presence of CD8 mRNA expression in few other species as in human^**37**^, in canine^**38**^ and in cows^**39**^. Preliminary study with CD4 gene expression in sheep had been studied in liver infected with cystic Echinococcus^**40,41**^. Apart from the role on bacterial, protozoal, parasitic infestation, CD4 and CD8 were observed to have a positive role in viral infection^**42**^, thus multiple roles of CD4 and CD8 were depicted^**43**^. Fetus specific T cell modulation was observed during fertilization, implantation and pregnancy^44^, with positive influence of CD4 and CD8 coreceptor gene expression^**45**^. Different molecular biology techniques were employed for studying CD4 and CD8 with differential mRNA expression profiling for Mycobacterium tuberculosis^46^. Advanced NGS studies were employed to assess gene expression profiling of sheep ovary^**47**^. We had earlier characterized CD14 gene in different species of livestock^**48, 49, 50**^. CD8 and CD4 expression profile revealed that these may be used as future marker for pregnancy detection in sheep, which always remain a challenge for sheep industry. The technique may even be standardized for detection of litter size, which is a economic trait and of great importance in sheep which is prolific.

## Materials and methods

### Sample

To undertake the present investigations, apparently healthy six pregnant (gravid) and six non-pregnant (non-gravid) uterus of sheep were procured from the Haringhata meat slaughter unit, Nadia, Govt. of West Bengal, India. The samples were harvested immediately after slaughter and taken into the ice box after necessary cleaning. Tissues were collected from different parts of uterus and preserved in 10% neutral buffered formalin (NBF) for histological and immunohistochemical studies.

Simultaneously, tissue samples from different segment of uterus (middle horn uterus and tip of the horn) of pregnant and non-pregnant ewes were collected into 5 ml eppendorf tubes separately with trizol (Ambion, Cat.No. 155960) and preserved at 4°C for initial 24 hrs and subsequently at −20°C till estimation for the mRNA expression.

### Real time PCR

Total RNA was estimated from the middle horn of uterus and tip of the horn of both the gravid and non-gravid uterus by Trizol method^**48,49,50,51**^ and quantitative analysis of total RNA were performed using formaldehyde gel electrophoresis. The 28S rRNA and 18Sr RNAwere demarcated the quality of RNA.First strand cDNA was synthesised by the process of reverse transcriptase polymerase chain reaction (rt-PCR) in the automated temperature maintained thermocycler mechine. M-MLVRT (200 u/μl) was used as reverse transcriptase enzyme.All the primers were designed using primer 3 software (v. 0. 4.0) as per the recommended criteria. The primers used are listed in Table1. Equal amount of RNA (quantified by Qubit fluorometer, Invitrogen), wherever applicable, were used for cDNA preparation (Superscript III cDNA synthesis kit; Invitrogen). All qRT-PCR reactions were conducted on ABI 7500 fast system. Each reaction consisted of 2 μl cDNA template, 5 μl of 2X SYBR Green PCR Master Mix, 0.25 μl each of forward and reverse primers (10 pmol/μl) and nuclease free water for a final volume of 10 μl. Each sample was run in duplicate. Analysis of real-time PCR (qRT-PCR) was performed by delta-delta-Ct (ΔΔCt) method.

**Table 1:**
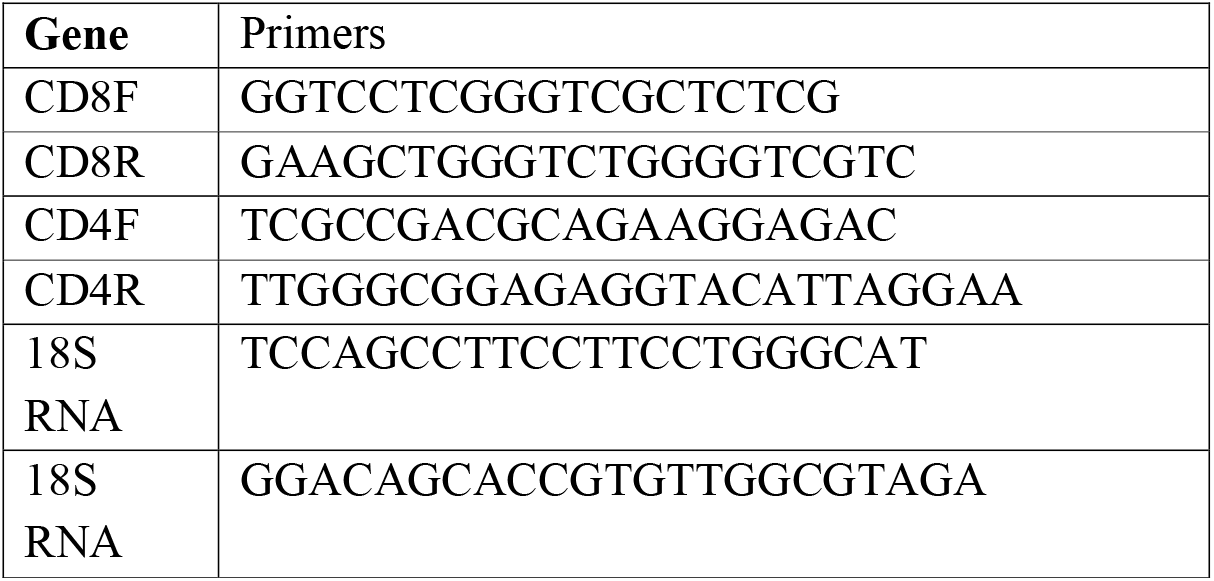
List of primers involves in Real time PCR of ovine cDNA

The entire reactions were performed in triplicate (as per MIQE Guidelines) and experiment repeated twice, in 20μl reaction volume, using FastStart Essential DNA Green Master (Himedia) on ABI 7500 system.

### Immunohistochemistry

Paraffin tissue blocks were prepared by standard manual alcohol-acetone protocol. The 5-6 μm thick paraffin sections obtained from different segments of the uterus (middlehorn of uterus and tip of the horn) of both the pregnant and non-pregnant ewes, were taken on Millennia 2.0 adhesion slides (Cat. No. 71863-01, abcam).The tissue sections were de-paraffinized and hydrated in distilled water. The tissues slides were covered with trypsin enzymatic antigen retrieval solution (Cat. No. ab970, abcam) and kept in incubator in humid environment at 37°C for 5-10 minutes. The sections were then incubated for 60 minutes in peroxidase blocking solution (Lot. No. 00065614, Dako) at room temperature to block non-specific antibody binding activity. After subsequent washing with phosphate buffer saline (PBS), the sections were incubated at 37°C for 2 hours in humid environment with mouse monoclonal anti CD8 antibody (Cat. No. MA1-80900, Thermo Fisher Scientific) and CD4 antibody (Mouse anti Sheep CD4 antibody Cat no. MCA2213GA, abCam) in 1: 200 dilution. Immunoreactivity was detected after one hour incubation at 37°C with secondary antibody, Rabbit anti-mouse IgG H&L (HRP Conjugated, Cat. No. ab6728; abcam) in dilution 1:200. Slides were then rinsed 3 times in PBS for 5 minutes each, followed by treatment with freshly prepared DAB solution for 3 minutes (DAB substrate, Cat. No. 34001, Thermo Fisher Scientific). The sections were counter stained with Mayer’s haematoxylin, hydrated in ethanol, cleared in xylene and then mounted in DPX^**52**^.

### Statistical analysis

Researchers count the percentage of positive immune labeled cells over the total cells in each selected area. This method can be automated with the use of special plugins for computer counting of general amount of cells and positively stained cells in case of immunohistochemistry.

Descriptive statistics with mean and standard error were estimated through SYSTAT package for the expression level analyzed through real time PCR and presented accordingly in graph.

## Conclusion

The lymphocyte population was more at the middle horn of the uterus as compared to the tip of the uterus in both gravid and non-gravid ewes. The occurrence of CD8+ cells were found within the luminal and glandular epithelium of the caruncular and inter caruncular areas. In gravid uterus the population of CD8+ T cell was comparatively more. The middle horn of the uterus revealed maximum expression in both the gravid and non-gravid ewes. The current observations demonstrate that during pregnancy in ewe the immune system responds to changes in the maternal environment to maintain the size and function of the CD8 T-cell compartment, as revealed by immunohistochemistry studies. Studies at genomic level revealed an increased mRNA expression of CD8 and CD4 gene, which is the first report, confirmed the above findings. CD8 expression pattern was higher in middle part of the pregnant uterus with both the techniques studied, indicating the importance of CD8 in case of recurrent abortion in non-infective condition.CD8 expression was higher in gravid uterus compared to that of CD4 gene expression. CD8 and CD4 may be effectively used as marker for pregnancy detection in sheep.

## Supporting information

Highlights

## Acknowledgement

The authors are very much grateful to Vice-Chancellor, West Bengal University of Animal and Fishery Sciences; Dean, Faculty of Veterinary & Animal Sciences, WBUAFS and Head of the Department of Veterinary Anatomy and Histology, WBUAFS, Kolkata, India for providing necessary facilities to investigate the present study. The authors greatly acknowledge the funding provided by Department of Biotechology, Ministry of Science and Technology, Govt. of India (Grant no. BT/Bio-CARe/04/10100/2013-14).

## Author Contribution Statement

MI, AP and PD has planned and designed the research. MI, SB and AP had conducted the research. AP, PD and MI had analyzed the data. MI, AP and PD had drafted the manuscript and prepared the figures. SB had critically reviewed the article.

## Reference

1. Tizard, I. R.. An introduction to Veterinary Immunology. 8^th^ edn, WB Saunders. (1977)

2. Norton M. T., Fortner K. A., Oppenheimer K. H. & Bonney E. A. Evidence that CD8 T-cell homeostasis and function remain intact during murine pregnancy. Immunology, 131(3), 426–437 (2010).

3. Miceli M. C. & Parnes J. R. The roles of CD4 and CD8 in T cell activation. Seminars in Immunology. May, 3(3), 133–41 (1991).

4. Artyomov M. N., Lis M., Devadas S., Davis M. M. & Chakraborty A. K. CD4 and CD8 binding to MHC molecules primarily acts to enhance Lck delivery. Proceedings of the National Academy of Sciences. 107(39), 16916–16921 (2010).

5. Green A. M., DiFazio R. & Flynn J. L. IFN-γ from CD4 T cells is essential for host survival and enhances CD8 T cell function during Mycobacterium tuberculosis infection. The Journal of Immunology. Jan 1, 190(1), 270–277 (2013).

6. Engwerda, C. R., Ng, S. S. & Bunn, P. T. The regulation of CD4+ T cell responses during protozoan infections. Frontiers in immunology. Oct 13, 5, 498 (2014).

7. McCune J. M. The dynamics of CD4+ T-cell depletion in HIV disease. Nature, Apr, 410(6831), 974–979 (2001).

8. Vidya Vijayan K. K., Karthigeyan K. P., Tripathi S. P. & Hanna L. E. Pathophysiology of CD4+ T-cell depletion in HIV-1 and HIV-2 infections. Frontiers in immunology. May 23, 8, 580 (2017).

9. Pal A., Chatterjee P.N., Das S., Batobyal S., Biswas P. & Sharma A. Biodiversity among sheep and goat reared under different agroclimatic regions of West Bengal, India. Indian Journal of Animal Science. 87(1), 80–86 (2017).

10. Sahana G., Gupta S. C. & Nivsarkar A. E. Garole: The prolific sheep of India. AnimalGeneticResources/Resourcesgénétiquesanimales/Recursosgenéticosanimales. 31, 55–63 (2001).

11. Pradhan M., Pal A., Samanta A.K., Banerjee S. & Samanta R. Mutations in cytochrome B gene effects female reproduction of Ghungroo pig. Theriogenology. 119, 121–130 (2018).

12. Entrican G. & Wheelhouse N. M. Immunity in the female sheep reproductive tract. Veterinary research. 37(3), 295–309 (2006).

13. Wira C. R., Rodriguez-Garcia M., & Patel M. V. The role of sex hormones in immune protection of the female reproductive tract. Nature reviews Immunology. Apr, 15(4), 217 (2015).

14. Shanmugasundaram U., Critchfield J. W., Pannell J., Perry J., Giudice L. C., Smith□McCune K., Greenblatt R. M. & Shacklett B. L. Phenotype and Functionality of CD 4+ and CD 8+ T Cells in the Upper Reproductive Tract of Healthy Premenopausal Women. American journal of reproductive immunology. Feb, 71(2), 95–108 (2014)

15. Sathaliyawala T., Kubota M., Yudanin N., Turner D., Camp P., Thome J. J. & Kato T. Distribution and compartmentalization of human circulating and tissue-resident memory T cell subsets. Immunity. Jan 24, 38(1), 187–197 (2013).

16. Gottshall S. L. & Hansen P. J. Regulation of leucocyte subpopulations in the sheep endometrium by progesterone. Immunology. 76(4), 636 (1992).

17. Lee C. S., Gogolin-Ewens K. & Brandon M. R. Identification of a unique lymphocyte subpopulation in the sheep uterus. Immunology. 63(1), 157 (1988).

18. Hansen P. J. & Liu W. J. Immunological aspects of pregnancy: concepts and speculations using the sheep as a model. Animal Reproduction Science. 42(1-4), 483–493 (1996).

19. Meeusen E., Fox A., Brandon M. & Lee C. S. Activation of uterine intraepithelial γδ T cell receptor positive lymphocytes during pregnancy. European journal of immunology. 23(5), 1112–1117 (1993).

20. Cobb S. P. & Watson E. D. Immunohistochemical study of immune cells in the bovine endometrium at different stages of the oestrous cycle. Research in veterinary science. 59(3), 238–241 (1995).

21. Kaeoket K., Dalin A. M., Magnusson U. & Persson E. The sow endometrium at different stages of the oestrous cycle: studies on the distribution of CD2, CD4, CD8 and MHC class II expressing cells. Animal reproduction science. 68(1-2), 99–109 (2001).

22. Perez-Martinez M., Luna J., Mena R. & Romano M. C. Lymphocytes and T lymphocyte subsets are regionally distributed in the female goat reproductive tract: influence of the stage of the oestrous cycle. Research in veterinary science. 72(2), 115–121 (2002).

23. Majewski A. C., Tekin S. & Hansen P. J. Local versus systemic control of numbers of endometrial T cells during pregnancy in sheep. Immunology. 102(3), 317–322 (2001).

24. Bischof R. J., Brandon M. R. & Lee C. S. Cellular immune responses in the pig uterus during pregnancy. Journal of reproductive immunology. 29(2), 161–178 (1995).

25. Gogolin-Ewens K. J., Lee C. S., Mercer W. R. & Brandon M. R. Site-directed differences in the immune response to the fetus. Immunology. 66(2), 312 (1989).

26. Tabibzadeh S. Proliferative activity of lymphoid cells in human endometrium throughout the menstrual cycle. The Journal of Clinical Endocrinology & Metabolism. 70(2), 437–443 (1990).

27. Akbalik M. E., Liman N., Sagsoz H. & GuneySaruhan B. Tissue distribution of some immune cells in bovine reproductive tract during follicular and luteal phase. Microscopy research and technique. 81(3), 315–331 (2018).

28. Vasudevan S., Kamat M. M., Walusimbi S. S., Pate J. L. & Ott T. L. Effects of early pregnancy on uterine lymphocytes and endometrial expression of immune-regulatory molecules in dairy heifers. Biology of reproduction. 97(1), 104–118 (2017).

29. Lissauer D., Piper K., Goodyear O., Kilby M. D. & Moss P. A. Fetal-specific CD8+ cytotoxic T cell responses develop during normal human pregnancy and exhibit broad functional capacity. The Journal of Immunology. 189(2), 1072–1080 (2012).

30. Guerin L. R., Prins J. R. & Robertson S. A. Regulatory T cells and immune tolerance in pregnancy: a new target for infertility treatment? Hum Reprod Update, 15, 517–35 (2009).

31. Tilburgs T., Scherjon S. A., Vander Mast B. J. & Haadnoot G. W., Versteeg-vander Voort-Maarschalk, M., Roelen, D.I. Fetal maternal HCA-C mismatch is associated with decidual T cell activation and induction of functional T regulatory cells. J. of Reproduction Immunology. 82, 148–157 (2009).

32. Head J. R. & Billingham R. E. Concerning the immunology of the uterus. Am. J. reprod. Immunol. 10, 76 (1986).

33. Quayle A. J. The innate and early immune response to pathogen challenge in the female genital tract and the pivotal role of epithelial cells. Journal of reproductive immunology, 57(1-2), 61–79 (2002).

34. Watnick A. S. & Russo R. A. Survival of skin homografts in uteri of pregnant and progesterone-estrogen treated rats. Proceedings of the Society for Experimental Biology and Medicine. 128(1), 1–4 (1968).

35. Hansen P. J. & Liu W. J. Immunological aspects of pregnancy: concepts and speculations using the sheep as a model. Animal Reproduction Science. 42(1-4), 483–493 (1996).

36. Clark D. A., Arck P. C. & Chaouat G. Why did your mother reject you? Immunogenetic determinants of the response to environmental selective pressure expressed at the uterine level. American Journal of Reproductive Immunology. 41(1), 5–22 (1999).

37. Diaz L. S., Stone M. R., Mackewicz C. E. & Levy J. A. Differential gene expression in CD8+ cells exhibiting noncytotoxic anti-HIV activity. Virology. 311(2), 400–409 (2003).

38. Schäfer S., Beceriklisoy H. B., Budik S., Kanca H., Aksoy O. A., Polat B. & Aslan S. Expression of Genes in the Canine Pre-implantation Uterus and Embryo: Implications for an Active Role of the Embryo Before and During Invasion. Reproduction in domestic animals. 43(6), 656–663 (2008).

39. Barański W., Kaleczyc J., Zduńczyk S., Podlasz W., Długołęcka-Malinowska E. & Janowski T. Distribution of CD 14+ macrophages, CD4+, CD8+ lymphocytes and mRNA expression of inducible nitric oxide synthase in the endometrium of repeat breeding cows. Polish journal of veterinary sciences, 16(3), 443–451 (2013).

40. Hui W., Jiang S., Liu X., Ban Q., Chen S. & Jia B. Gene expression profile in the liver of sheep infected with cystic echinococcosis. PloS one. Jul 28, 11(7), e0160000 (2016).

41. Bush S. J., McCulloch M. E. B., Muriuki C., Salavati M., Davis G. M., Iseabail L. Farquhar I. L., Lisowski Z. M., Archibald A. L., Hume D. A. & Clark E. L. Comprehensive Transcriptional Profiling of the Gastrointestinal Tract of Ruminants from Birth to Adulthood Reveals Strong Developmental Stage Specific Gene Expression. G3: Genes, Genomes, Genetics. Feb 1, 9(2), 359–373 (2019).

42. Nascimbeni M., Shin E. C., Chiriboga L., Kleiner D. E. & Rehermann B. Peripheral CD4+ CD8+ T cells are differentiated effector memory cells with antiviral functions. Blood. Jul 15, 104(2), 478–486 (2004).

43. Scherjon S., Lashley L., Van Der Hoorn, M. L. & Claas F. Fetus specific T cell modulation during fertilization, implantation and pregnancy. Placenta. 32, S291–S297 (2011).

44. Ellmeier W., Sawada S. & Littman D. R. The regulation of CD4 and CD8 coreceptor gene expression during T cell development. Annu. Rev. Immunol. 1999. 17, 523–54 (1999).

45. Koretzky G. A. Multiple roles of CD4 and CD8 in T cell activation. The journal of immunology. Sep 1, 185(5), 2643–2644 (2010).

46. Chen H. Y., Shen H., Jia B., Zhang Y. S., Wang X. H. & Zeng X. C. Differential gene expression in ovaries of Qira black sheep and Hetian sheep using RNA-Seq technique. PloS one. Mar 19, 10(3), e0120170 (2015).

47. Cliff J. M., Andrade I. N., Mistry R., Clayton C. L., Lennon M. G., Lewis A. P. & Dockrell H. M. Differential gene expression identifies novel markers of CD4+ and CD8+ T cell activation following stimulation by Mycobacterium tuberculosis. The Journal of Immunology. Jul 1, 173(1), 485–493 (2004).

48. Pal A., Sharma A., Bhattacharya T. K., Chatterjee P. N. & Chakravarty A. K. Molecular characterization and SNP detection of CD 14 gene in crossbred cattle. Molecular Biology International. vol. 2011, Article ID 507346, 13 pages, 2011. doi:10.4061/2011/507346. (2011).

49. Pal, A. and Chatterjee, P.N. 2009. Molecular cloning and characterization of CD14 gene in goat. J. of Small Ruminant Research. 82: 84–87.

50. Pal, A., Chatterjee, P.N., and Sharma, A. 2014. Sequence characterization and polymorphism detection in bubaline CD14 gene. Buffalo Bulletin. 32:2, 138–156.

51. Pal, A, Pal, A. Jr., Mallick, A.I., Biswas, P. And Chatterjee, P.N. 2019. _Molecular characterization of Bu-1 and TLR2 gene in Haringhata Black chicken. Genomics. 112(1):472–483. https://doi.org/10.1016/j.ygeno.2019.03.010

52. Pal, A., Pal, Ab and Chakravarty, A.K. 2020. Mutations in Growth hormone gene affect stability of protein structure leading to reduced growth, reproduction and milk production in Cross Bred cattle-an insight. Domestic Animal Endocrinology. https://doi.org/10.1016/j.domaniend.2019.106405

