## Supplementary material for "Better expression profile of CD8 and CD4 gene in uterus of pregnant ewe in comparison to non-pregnant - a novel report": Highlights

- CD8 and CD4 gene expression was better in pregnant uterus of sheep
- Differential mRNA expression profile for above genes are reported for the first time in sheep
- Immunohistochemistry confirms the above findings
- CD8 and CD4 expression may be regarded as the useful factor for maintenance of pregnancy.
